# Revisiting colorectal cancer tumorigenesis with spatially-resolved gene expression profiling

**DOI:** 10.1101/2021.09.30.462502

**Authors:** Jessica Roelands, Manon van der Ploeg, Hao Dang, Jurjen J. Boonstra, James C.H. Hardwick, Lukas J.A.C. Hawinkels, Hans Morreau, Noel F.C.C. de Miranda

## Abstract

Early detection and treatment are paramount to the clinical outcome of patients with colorectal cancer (CRC). Deciphering the dynamic interactions that occur between epithelial cells and stromal cells during tumorigenesis requires in-depth analyses of early-stage CRC lesions in spatial context. Here we employed spatially-resolved gene expression profiling to dissect molecular processes that associate with malignant transformation in CRC. We provide the transcriptional landscapes of colorectal cancer tumorigenesis from healthy mucosa, through different degrees of dysplasia, to cancer. The complementary examination of epithelial and stromal fractions allowed us to define whether specific oncogenic processes involved cancer cells, stromal cells, or the tumor microenvironment as a whole. We identified several genes that were consistently deregulated during CRC onset that could serve as clinical biomarkers for early-stage CRC. Furthermore, we uncovered an essential role for the innate immune system during CRC tumorigenesis.

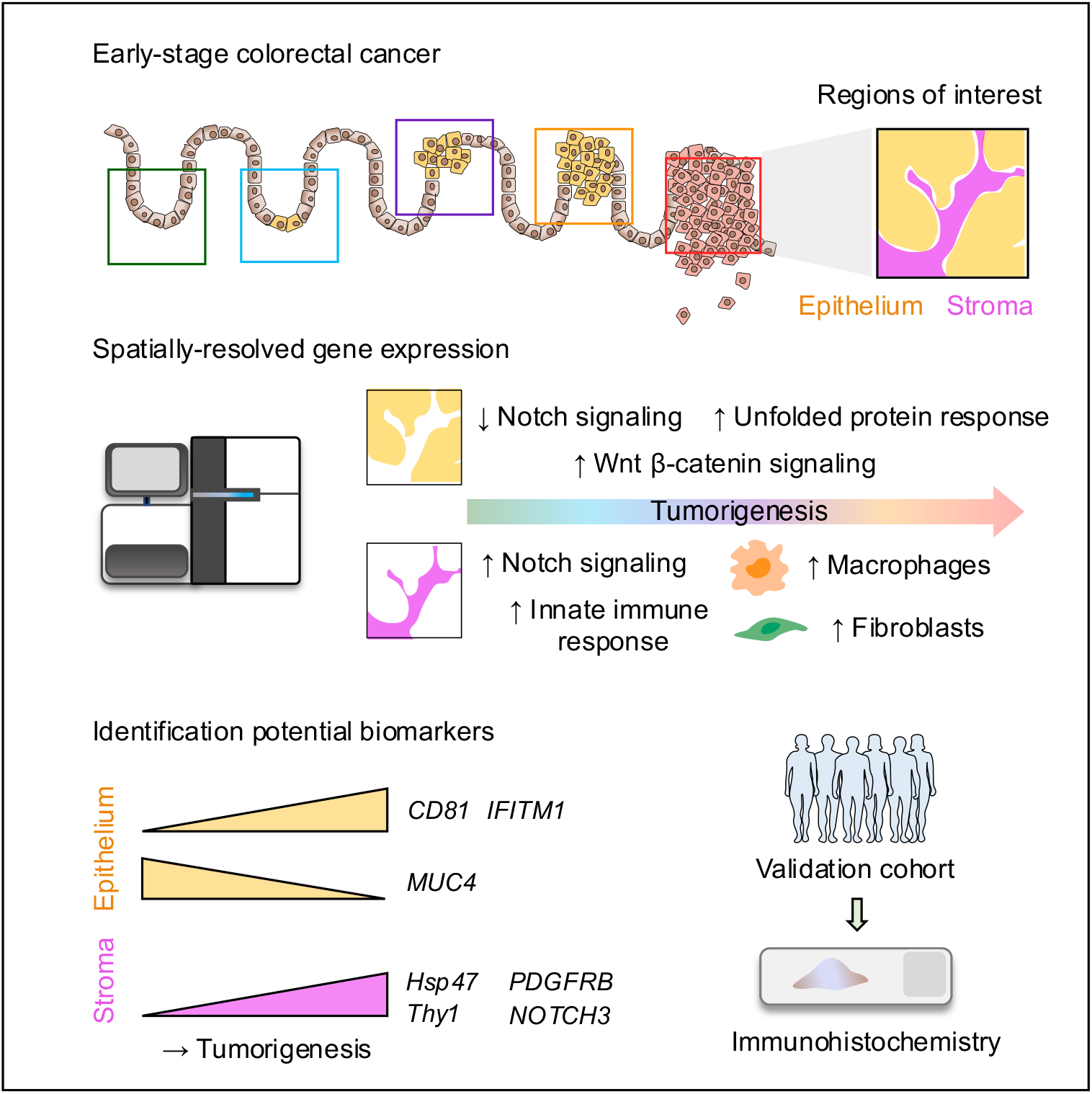

## Introduction

Colorectal cancers (CRCs) arise as a consequence of the gradual accumulation of (epi-) genetic alterations in epithelial cells in the colon and rectum, ultimately leading to uncontrolled proliferation and malignant transformation^1^. The paradigm of the adenoma-carcinoma sequence has typically been centred at cancer cells; however, it is now clear that cancer progression involves heterologous interactions between aberrant cells and their surrounding microenvironment^2^. Thus far, studies that investigated the role of the microenvironment in early stages of CRC have been relatively scarce. This is partly due to the (previously) limited availability of technological approaches for in-depth investigation of tumorigenic processes in spatial context.

The process of CRC tumorigenesis, from benign to malignant lesions, typically takes years or even decades^1,3,4^. From a clinical point of view, this represents a suitable time-window for early disease interception. Population screening programs aim at the early diagnosis and treatment of CRC, hereby reducing patient morbidity and mortality^5^. These programs have resulted in an increased number of patients diagnosed with early-stage CRC (e.g., pT1 CRC) that are amenable to less invasive clinical interventions that include endoscopic removal of tumours. The dilemma whether conservative (e.g., endoscopic) procedures should be followed by extensive curative procedures (e.g., surgical resection) in pT1-staged CRCs, is generally addressed by morphologic and histologic evaluation of the lesions^6,7^. However, this strategy is suboptimal, as reflected by substantial over-treatment of patients (∼80-90%)^8,9^. Therefore, it is crucial to define biomarkers that reliably discern patients with aggressive lesions at risk of metastatic disease. To obtain high-quality biomarkers that can by employed in the clinic, it is imperative to decipher the biological processes at the basis of cancer initiation.

In recent years, technologies for spatially resolved gene expression profiling have evolved rapidly, offering opportunities to define transcriptomic changes in spatial context^10^. One of these technologies, the NanoString GeoMx digital spatial profiling (DSP) assay, employs RNA probes to quantify the abundance of transcripts in formalin-fixed paraffin-embedded (FFPE) tissue sections^11^. Transcript abundance can be measured in defined regions of interest in the tissue, thereby allowing the separate interrogation of cellular compartments within the same tissue. To elucidate the biological processes that occur during malignant transformation in CRC, we interrogated the transcriptional landscapes of normal mucosa, low-, and high-grade dysplasia, and cancer in eight pT1 CRC lesions.

This work provides unique biological insights into CRC tumorigenesis through the application of transcriptomic spatial profiling. We identified and validated the expression of several genes of interest that were consistently deregulated during CRC onset. These targets could potentially serve as novel biomarkers and targets for early interception and prevention of malignant transformation. Furthermore, an essential role for the innate immune system during early stages of CRC development was pinpointed by our approach. Importantly, the parallel analysis of epithelial and stromal fractions provided us the opportunity to define whether biological alterations during CRC tumorigenesis are driven by cancer cells, stromal cells, or the complete tumor microenvironment.

## Results

### Spatially resolved gene expression profiling of the transcriptomic landscape of CRC tumorigenesis

To decipher the molecular changes that occur during the onset and progression of CRC, we profiled eight pT1 CRC samples using GeoMx DSP technology. For each sample, nine ROIs were selected, encompassing normal mucosa, areas of transition between normal and dysplasia, low– and high grade dysplasia, and carcinoma (**Fig. 1a**). To specifically examine transcriptional alterations within epithelial and stromal compartments, we interrogated, separately, cytokeratin positive (PanCK+) and vimentin positive (Vimentin+) fractions within each ROI (**Fig. 1b**).

**Fig. 1.**
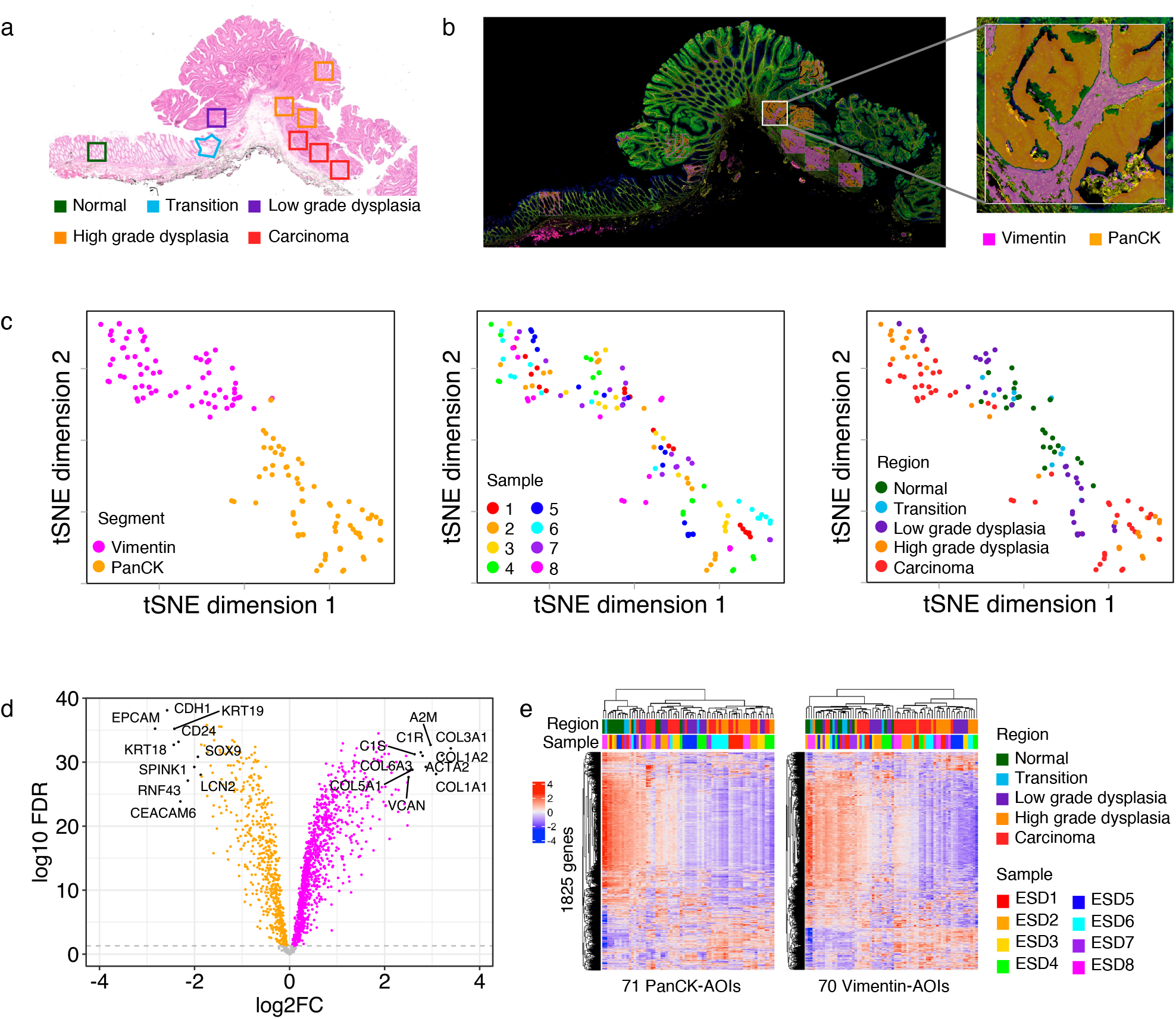
Transcriptional alterations in early-stage CRC defined by Digital Spatial Profiling. (**a**) Representative example of pT1 CRC sample stained by H&E. Selected ROIs with different histological status are annotated. (**b**) Immunofluorescent image of the same sample as in panel (**a**), stained with fluorescent antibodies against PanCK and Vimentin. Artificial overlay of implemented segmentation is indicated for each ROI, visualizing Vimentin+ (pink) and PanCK+ (orange) segments. Inset: higher magnification of an individual ROI (high grade dysplasia). (**c**) Dimension reduction of the expression of all quantified genes per AOI by tSNE. tSNE plots are annotated by segment (left), sample ID (middle), and histological region (right). Each datapoint reflects a single AOI. (**d**) Volcano plot of differentially expressed genes between Vimentin and PanCK segments by paired t-test. FDR is calculated using the Benjamini-Hochberg method. (**e**) Heatmaps of expression for all genes (n=1825) using unsupervised clustering for all PanCK AOIs (n= 71, left) and all Vimentin AOIs (n=70, right). Heatmaps are annotated by histological region and sample ID.

Data exploration using dimension reduction by t-Distributed Stochastic Neighbor Embedding (tSNE) revealed, as expected, a clear segregation of the AOIs by segment (PanCK/Vimentin) (**Fig. 1c**, *left panel*). The large majority of genes in the Cancer Transcriptome Atlas (CTA) gene collection was differentially expressed between segments; 583 genes were upregulated in the epithelial segments and 1068 genes were upregulated in the stromal segments (paired t-test; Benjamini-Hochberg method FDR < 0.05) (**Fig. 1d**). Samples of individual patients did not form major clusters in the high-dimensional space (**Fig. 1c**, *middle panel*). Instead, mapping of AOIs in the tSNE was related to the histologic features of the profiled regions (**Fig. 1c**, *right panel*). Interestingly, normal and transition areas separated from high grade dysplasia and carcinoma AOIs within both the epithelial and the stromal segments. These findings indicate that CRC onset is associated with transcriptional alterations in the (pre-) malignant epithelial cells themselves as well as the surrounding stromal cells. Inter-patient variability was most noticeable in epithelial AOIs of high grade dysplasia and carcinoma (**Fig. 1c**, *middle panel*) that also is likely to contain the highest (epi-) genetic heterogeneity.

For a high-level overview of the expression data, we subsequently visualized the expression of all 1,825 CTA genes in unsupervised clustered heatmaps, per segment (**Fig. 1e**). In total, 242 genes were increasingly expressed during tumorigenesis in the epithelial AOIs and 288 genes in the stromal AOIs (Spearman correlation, FDR < 0.05). Interestingly, a larger number of genes decreased during CRC onset (i.e., 1,152 genes in the epithelium and 857 genes in the stroma, Spearman correlation, FDR < 0.05). Genes with reduced expression during tumorigenesis demonstrated a highly consistent expression pattern across samples, whereas genes that were upregulated revealed more heterogeneous patterns (**Fig. 1e, Extended Data Fig. 1**). This observation likely reflects the common loss of physiological features of the colorectum during malignant progression across samples.

### Biological processes associated CRC onset are distinct between epithelium and stroma

We aimed to identify core molecular networks associated with tumorigenesis. All genes that were highly associated with histologic progression of disease (Spearman correlation coefficient > 0.5, FDR < 0.05) were included for core network analysis using Ingenuity Pathway Analysis^12^. This resulted in the identification of 86 genes associated with tumorigenesis in the epithelium and 131 genes in the stromal fraction. In the former, and unsurprisingly, the key network found to be activated was centred around β-catenin activity and its downstream target genes such as *MYC*^13^ (**Fig. 2a**). In the stromal fraction, the core network associated with cancer onset and progression consisted of different interacting pathways (**Fig. 2b**), including activation of TGF-β signaling and Notch signaling, increased transcription of collagen genes, and components of the complement system. The TGF-β pathway has been described as one of the major pathways driving CRC tumorigenesis, in particular during the transition from adenoma to carcinoma^14,15^. The specific upregulation in the tumor microenvironment instead of the epithelial cells themselves, could be a consequence of the interaction of tumor cells with resident fibroblasts, which in turn leads to hyperactivation of TGF-β signaling in cancer-associated fibroblasts (CAFs)^16–19^. Notch signaling has also been described as a crucial signaling pathway for homeostasis of intestinal epithelial cells^20,21^, but also plays roles in shaping of the tumor vasculature, regulation of anti-tumor immune responses, and fibroblast activity^22^. On the other hand, the role of complement activation, a central component of innate immune defense, has more recently emerged as modulator of cancer progression, although its effects seem context-dependent^23^.

**Fig. 2.**
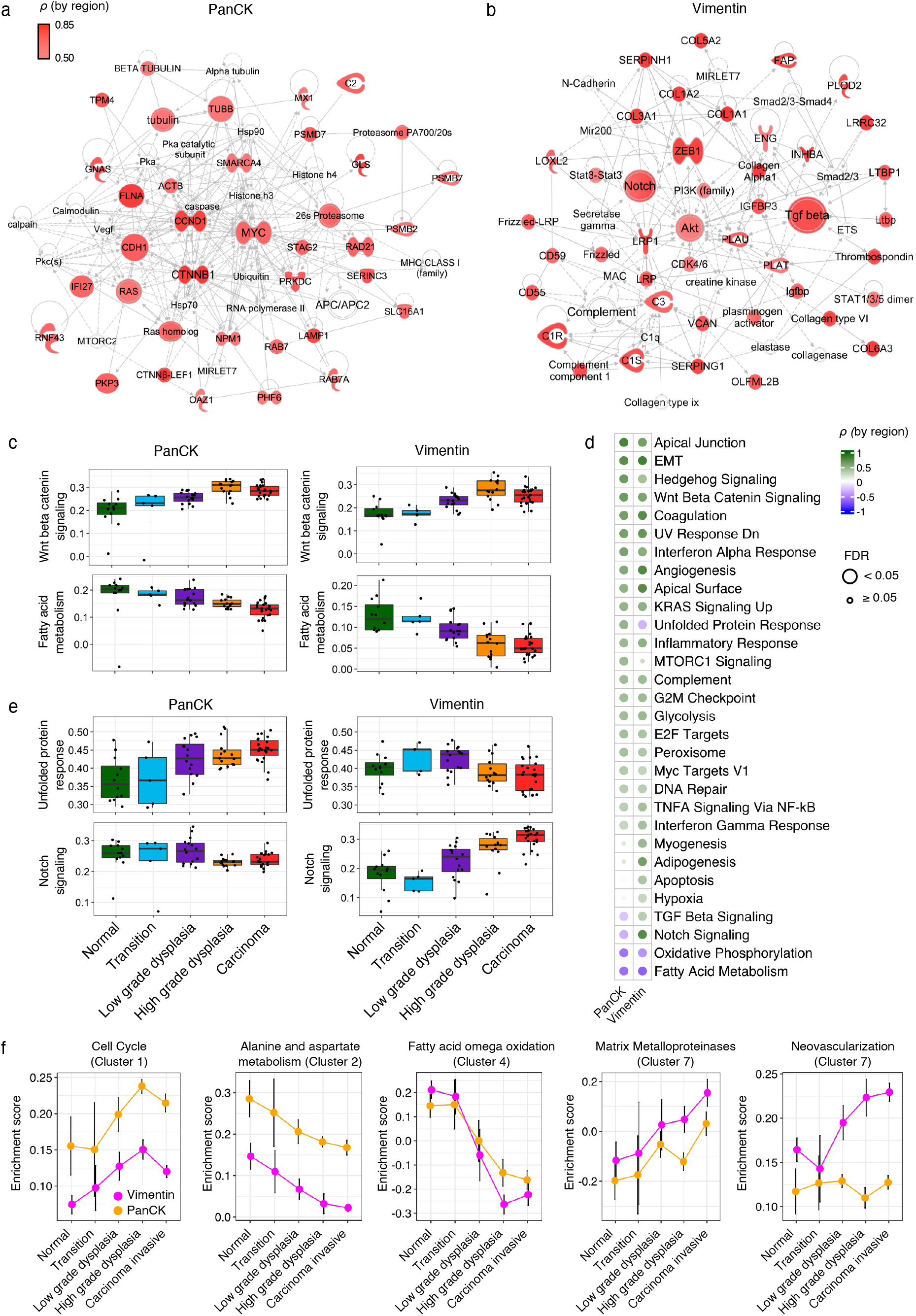
Key transcriptional networks and biological pathways associated with advancing histology in epithelial and stromal compartment. (**a, b**) Core Molecular Network that is associated with advancing histology identified by IPA. All genes that were associated with advancing histology with a Spearman correlation coefficient > 0.5 were used as input for IPA. This analysis was performed separately for PanCK+ (**a**) and Vimentin+ AOIs (**b**). (**c, e**) Boxplots of enrichment scores for cancer hallmark gene sets calculated by ssGSEA (y-axis) by histology (x-axis). Plots are separated by PanCK (left) and Vimentin (right) segments. (**d**) Heatmap of Spearman’s rho between enrichment scores for cancer hallmark pathways and histology as ordinal variable. Only pathways that are significantly associated with histology in either PanCK or Vimentin segments are shown. FDR is calculated using the Benjamini-Hochberg method. (**f**) Line graphs of highlighted pathways that are deregulated from normal tissue to carcinoma. Mean enrichment score and corresponding 95%-CI are indicated.

We subsequently interrogated alterations in cancer hallmark pathways^24^ between distinct histologic regions. As expected, most pathways that describe hallmarks of cancer, gradually increased their activity from normal tissue to carcinoma (**Fig. 2c**, *upper* panels, **Fig. 2d**). Two exceptions were noted that had an inverse association with malignant transformation (**Fig. 2d**), including oxidative phosphorylation and fatty acid metabolism (**Fig. 2c**, *lower panels*). The decrease of enrichment of genes associated with oxidative phosphorylation and concurrent increase in enrichment score of glycolysis pathway, points toward the switch to aerobic glycolysis^25,26^. While an increased fatty acid metabolism has been described as one of the key events in cancer metabolic reprogramming^27^, our results demonstrate a decrease in enrichment of fatty acid metabolism-related genes from normal tissue to carcinoma. This seemingly contradictory result most likely reflects the physiological role of the normal human colon in processing of fatty acids that is lost when the colonic tissue dedifferentiates.

Most cancer hallmark pathways demonstrated concordant alterations in the epithelial compartment and in the stromal compartment. However, some pathways were noted in which the directionality of the alterations was inverted in the epithelium versus the stromal compartment (**Fig. 2e**). For example, the enrichment of unfolded protein response pathway genes increased in the epithelium during tumor progression, whereas a decrease was observed in the stroma. Similarly, the enrichment of Notch pathway genes increased in the Vimentin segment, while a slight decrease was observed in the PanCK segment. These findings illustrate the relevance of the segregation of tissues in its major constituents, the epithelial and surrounding stroma and highlight that biologically relevant processes related to tumor progression might not be identified when averaging the contribution of cellular compartments to transcriptional signatures.

For a more comprehensive assessment of alterations during progression from normal tissue to carcinoma, we extended our analysis by employing a more inclusive collection of pathways. The WikiPathways database^28^ was used to extract non-redundant biological, cancer associated pathways suitable for single sample Gene Set Enrichment Analysis (ssGSEA). Unsupervised clustering of the pathways revealed seven distinct clusters (C1-C7), with different degrees of association with malignant transformation. Three clusters (C1, C2, and C4) were highly enriched in the epithelial segment. C1 pathways included biological processes associated with proliferation and DNA damage repair, which consistently increased during CRC onset, the “Cell Cycle” gene signature is shown as representative example in **Fig. 2f**. The majority of pathways clustered in C2 were associated with mitochondrial function, and demonstrated a decrease during malignant transformation (**Fig. 2f**). Similarly, pathways in C4 decreased during CRC onset, and included the PPAR signalling pathway and Fatty Acid Omega Oxidation. These alterations were paralleled in the stroma, indicating similar adaptations in both biological compartments (**Extended Data Fig. 2**). The pathways in clusters C5, C6, and C7 were mainly enriched in the stromal compartment. The stroma of dysplastic and invasive carcinoma regions was particularly enriched for pathways in C7, including pathways associated with TGF-β receptor signaling, focal adhesion, matrix metalloproteinases, and neovascularisation (**Fig. 2f**). These results demonstrate marked transcriptomic alterations during tumorigenesis and the existence of biological processes related to malignancy that are specifically enriched in the epithelial and stromal compartments.

### Immune-related alterations during CRC progression

Intrigued by the potential involvement of the complement system during CRC onset, we subsequently focused on immune-related transcriptional alterations. First, we performed immune cell deconvolution to estimate the relative abundancies of specific cell subsets in each AOI. In the stromal compartment, we defined multiple immune cell subsets with different relative abundancies between histological regions (ANOVA; Benjamini-Hochberg method FDR < 0.05). The relative abundance of plasma cells, B cells, CD8 T cells, CD4 T cells, T regulatory cells (T regs), γδ T cells, and NK cells, decreased from normal tissue to carcinoma (**Fig. 3a**). On the other hand, the relative abundance of fibroblasts, endothelial cells, and macrophages increased (**Fig. 3a**). These results point to the development of an immune-suppressed microenvironment accompanying tumor progression, which is in line with the pronounced upregulation of TGF-β (**Fig. 2e**)^29–31^.

**Fig. 3.**
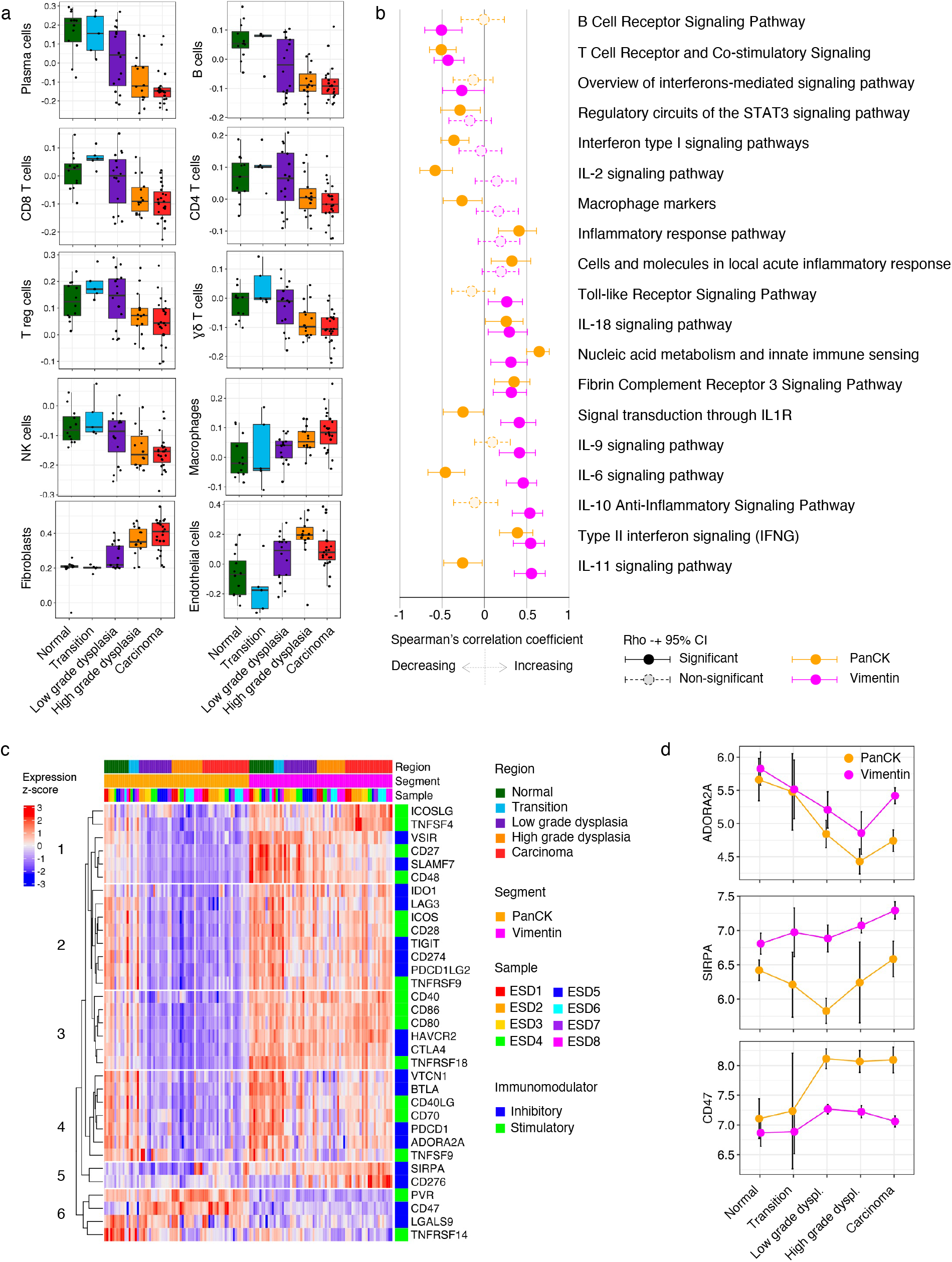
Immune-related perturbations in relation to CRC histology. (**a**) Boxplots of deconvoluted abundancies of distinct immune cell populations by histology of the ROI in the Vimentin segment. (**b**) Forest plot of Spearman’s Rho and corresponding 95% CI for the correlation between enrichment score of immune-related WikiPathway and histology as ordinal variable. Correlation was assessed in PanCK and Vimentin segments separately. (**c**) Expression of immunomodulators across distinct regions in the Vimentin segment. AOIs (columns) are ordered by histology and subsequently by sample ID. Genes (rows) are ordered by unsupervised clustering. Type of immunomodulator, either inhibitory or stimulatory, is indicated. (**d**) Line graphs of individual immunomodulators that are deregulated during tumorigenesis. Mean enrichment score and corresponding 95%-CI are indicated.

To define immune related alterations beyond the estimation of the cellular composition of the tumor microenvironment, we re-evaluated the altered enrichment of WikiPathways focusing on the immune-related gene sets. The majority of immune-related signatures (20/27) positively or negatively correlated with malignant transformation, reflecting substantial changes in the immune-microenvironment (**Fig. 3b**). Consistent with the decrease in deconvoluted B cells abundance, the enrichment scores of “B Cell Receptor Signaling Pathway” decreased from normal tissue to carcinoma. Other pathways that inversely correlated with advancing histology included “T Cell Receptor and Co-stimulatory Signaling” and “Overview of interferons-mediated signaling pathway”, reflecting a reduced number of T cells in the microenvironment. Different cytokine-related pathways were upregulated in the stroma during the progression from normal mucosa to carcinoma, including IFNG, IL-11, IL-10, IL-6, IL-9, IL1R, and IL-18 signaling pathways. Many of these cytokines have been described in the context of innate immunity (IL-1, IL-6, IL-10, IL-18, and TGF-β)^32^. These associations were infrequently mirrored by the epithelium. In fact, the IL-6, IL-11, and IL1R pathways actually had an inverse correction in the epithelium (**Fig. 3b**). Interestingly, we also identified two pathways linked to innate immunity that were consistently increased in both segments, including the “Fibrin Complement Receptor 3 Signaling Pathway” and “Nucleic acid metabolism and innate immune sensing” (see also **Extended Data Fig. 3b**). These results suggest the onset of an innate inflammatory response during CRC onset.

Next, we focused on immunomodulatory gene expression related to anti-tumor immunity. A large number of immunomodulators were increased in the stroma of normal and transition tissues (**Fig. 3c**), which likely reflects the tight balance that is in place in colonic tissues to provide immune defense but prevent exacerbated inflammation. Interestingly, a specific group of immunomodulatory genes revealed a biphasic pattern (cluster 4, **Fig. 3c**), with high expression in normal and transition tissue, decreased expression in dysplastic tissues, and slight increase in the stroma of carcinoma tissues. *ADORA2A* (coding adenosine A2A receptor) is shown as representative example in **Fig. 3d**. The “rescued” expression of these immunomodulatory genes in carcinoma tissues suggests the late adoption of mechanisms of T cell regulation during invasion of cancer cells to adjacent tissues. The immune immunomodulators *CD276* (B7-H3) and *SIRPA* (SIRPα) composed a distinct cluster (C5) and demonstrated increased expression in the stroma of high grade dysplasia and carcinoma. B7-H3 expression in stroma could be derived from myeloid cells but also from cancer-associated fibroblasts^33,34^ and lead to decreased anti-tumor activity by T cells^34,35^. SIRPα, or signal regulatory protein alpha, expression is mainly derived from myeloid cells^36^ which is in line with the observed increase in deconvoluted macrophages during tumor progression (**Fig. 3a**). Interestingly, the expression of its binding partner on tumor cells, *CD47*, was increased in the epithelial compartment in dysplastic and carcinoma tissues (**Fig. 3d**). Binding of CD47 on tumor cells to SIRPα on macrophages leads to the evasion of tumor destruction by inhibition of phagocytosis^37–40^. Our observations indicate that this “don’t eat me”-signal by tumor cells could be an effective mechanism of immune escape in these early-stage CRCs.

### Step-wise differential gene expression and biomarkers of CRC tumorigenesis

For a more in-depth analysis of the alterations during the progression from normal tissue to carcinoma, we assessed differential gene expression by step-wise comparisons between regions with distinct histologies (**Fig. 4a, b**). As expected, only small differences were identified between regions demonstrating morphological alterations between normal and dysplasia (transition) and normal tissue, both for the epithelial compartment as well as in the stroma (**Fig. 4a, b**, upper panels). However, these specific differentially expressed genes deserve some attention, as these potentially reflect the initial events of tumorigenesis. Interestingly, complement factor B (*CFB*) and C-C motif chemokine 20 (*CCL20*) were upregulated in the epithelium, while C-X-C Motif Chemokine Ligand 1 (*CXCL1*) was increased in the stroma of transition areas, providing signs of an early onset of inflammation and chemoattraction. Moving to areas of dysplasia, substantial differences in gene expression were observed, both in the epithelium as well as stromal compartments (**Fig. 4a, b**, lower panels).

**Fig. 4.**
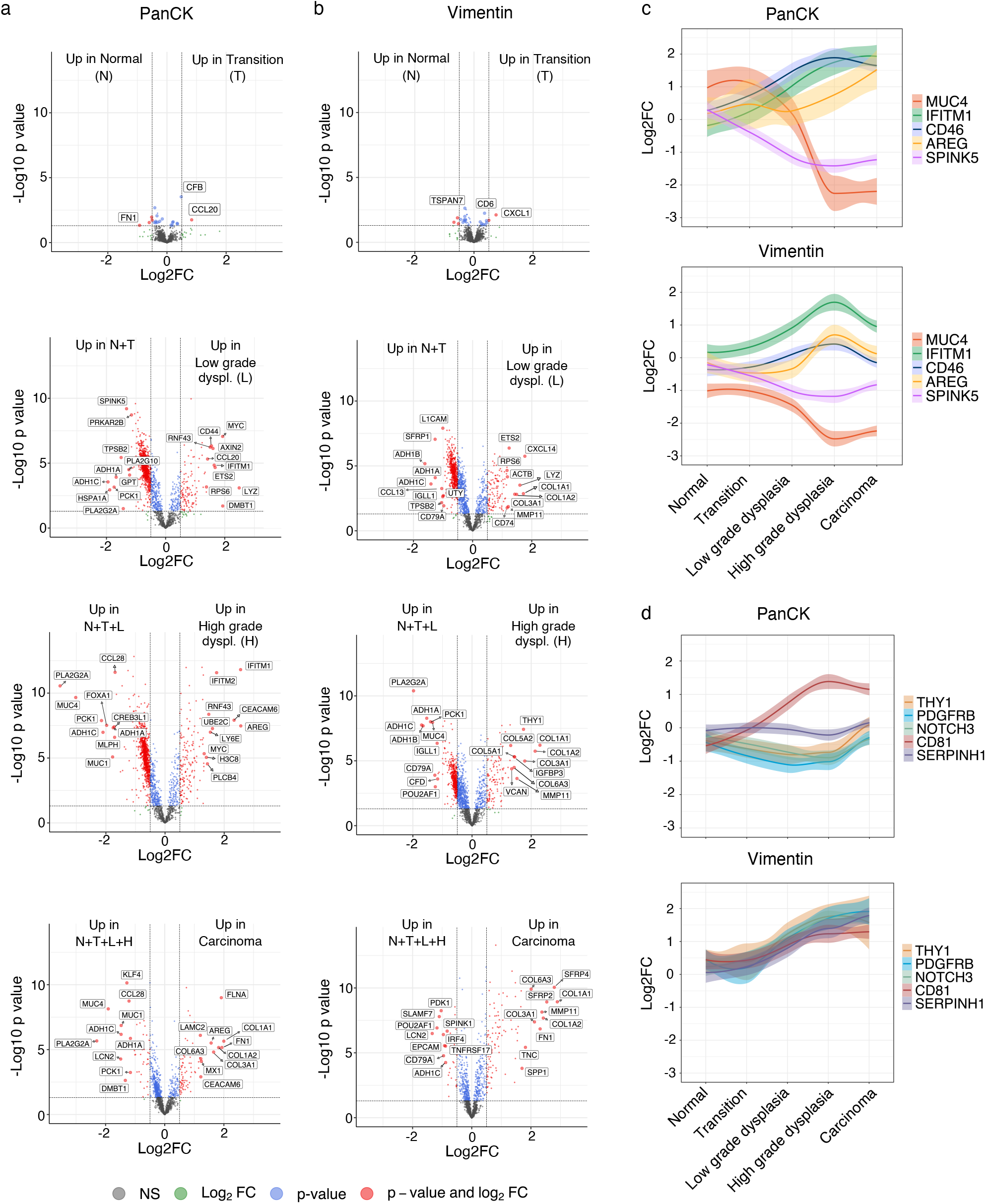
Step-wise differentially expressed genes from normal tissue to carcinoma in the epithelium and stroma. (**a, b**) Volcano plots showing differentially expressed genes between distinct histologies in step-wise comparisons (unpaired t-test). Reference group for comparisons are indicated. Analysis was performed for the epithelial compartment (PanCK) (**a**) and stroma compartment (Vimentin) (**b**), separately. Significance cutoff is a p value of 0.05 (-log10 p value of 1.30103) and log2FC cutoff is 0.5. Significant genes with the highest and lowest log2FC are labeled. (**c, d**) Selected genes of interest that demonstrate a clear and consistent step-wise alteration in gene expression from normal tissue to carcinoma. Genes were selected based on alterations in the PanCK segment (**c**) and in the Vimentin segment (**d**). The log2FC (y-axis) indicates the alteration as compared to the mean of all normal regions (PanCK and Vimentin combined). Line graph is generated using local polynomial regression fitting (loess-method) visualizing smoothed conditional means, corresponding confidence interval is indicated by the shadows.

Among the genes that were downregulated while progressing toward carcinoma in the PanCK segment were mucin genes *MUC4* and *MUC1*. In particular *MUC4* demonstrated a very consistent decrease in gene expression in regions of high grade dysplasia and carcinoma (**Fig. 4c**). Several additional genes were identified that were consistently altered from normal tissue to carcinoma. Based on the availability of commercial antibodies directed against the proteins encoded by these genes, ten genes of interest were selected. Beyond *MUC4*, selected genes with profound transcriptional alterations in the epithelium included *SPINK5, CD46, AREG, IFITM1*, and *CD81* (**Fig. 4c, d**). In addition, *THY1, PDGFRB, NOTCH3, CD81*, and *SERPINH1* were selected as candidates with a very consistent increase in the stroma (**Fig. 4d**).

Immunodetection of the MUC-4 protein in the eight samples of our spatially-resolved gene expression cohort, confirmed a clear epithelial expression in normal mucosa and transition areas, while minimal protein was detected in high grade dysplasia and carcinoma (**Fig. 5a, b**). An increase in expression of IFITM1, and CD81 was detected in the epithelium, also consistent with the observed alterations in gene expression (**Fig. 5a**). While we detected a higher abundance of the CD46 complement regulatory protein, the relative difference between normal and dysplastic tissues was not as pronounced as the changes observed in gene expression (**Fig. 4c**). For most of the selected genes of interest that were differentially expressed in the Vimentin segment, corresponding protein abundancies demonstrated concordant alterations in the stroma. For NOTCH3, Thy1, PDGFRB, and Hsp47, a clear upregulation of protein expression was observed along the step-wise progression from normal tissue to carcinoma. While gene expression of *NOTCH3* did not deviate substantially between regions of different histology in the PanCK segment (**Fig. 4d**), we did observe a decrease in protein abundance in the epithelium of these samples from normal tissue to carcinoma (**Fig. 5a**). For CD81, the observed increase in gene expression in the Vimentin segment did not translate into differential protein abundance in the stroma. We did not detect any protein expression Serine Peptidase Inhibitor Kazal Type 5 (*SPINK5*) and expression of Amphiregulin (*AREG*) could not reliably be assessed (data not shown).

**Fig. 5.**
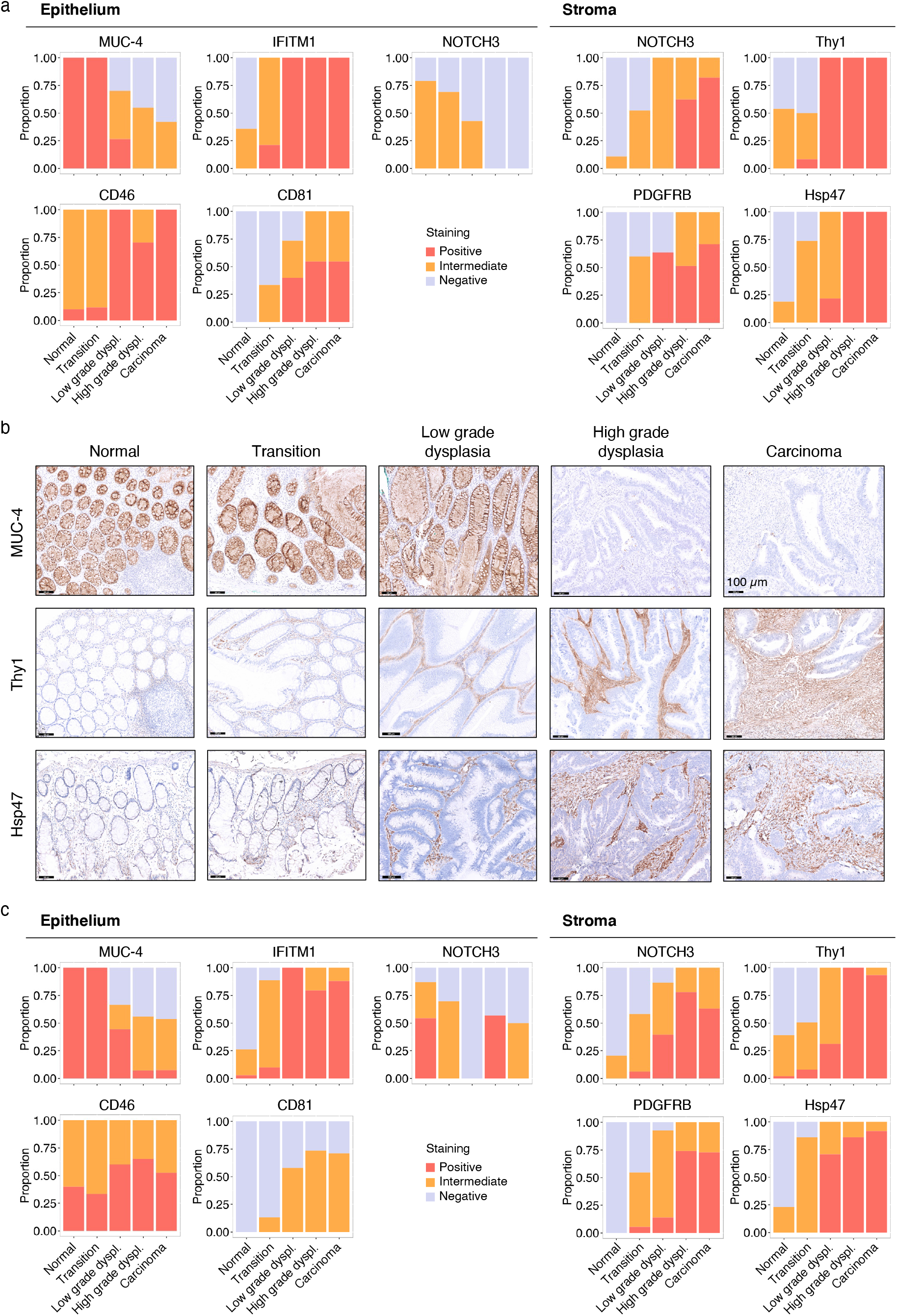
Differential protein abundance of candidate biomarkers in early-stage CRC. (**a**) Protein abundance of candidate biomarkers in the epithelium (*left*) and stroma (*right*) in the spatial transcriptomics early-stage CRC samples (n=8). (**b**) Representative images of IHC detection for MUC-4, Thy1, and Hsp47 across regions with distinct histology (normal, transition, low grade dysplasia, high grade dysplasia, and carcinoma). (**c**) Protein abundance of candidate biomarkers in independent validation set of 20 pT1 CRC samples. (**a, c**) Stacked bar chart reflects proportion of samples in each scored category.

To validate the differences in protein abundance, we assessed protein expression in an independent cohort of 20 pT1 CRC samples (**Fig. 5c**). We confirmed the downregulation of MUC-4, and upregulation of IFITM1, and CD81 in the epithelium during the transition of normal tissue to cancer. Of note, heterogeneous immunodetection patterns for MUC-4 and IFITM1 were observed within dysplastic and carcinoma tissues suggesting clonal heterogeneity. For CD46, only marginal differences were detected in the epithelium with a large variation between samples. For NOTCH3, protein abundance in the stroma increased, while a weak decrease was observed in the epithelium. These alterations in NOTCH3 protein expression parallels the changes in the overall enrichment score of the Notch signaling pathway (**Fig. 2e**). In the stroma, more consistent changes were detected for Thy1, PDGFRB, and Hsp47. Therefore, in particular these markers could be promising biomarkers for clinical utilization.

Overall, these results demonstrate that our methodological approach provides a rich source for identification of potential candidate biomarkers that are altered during the step-wise progression of cancer. Variability between patients is highest for candidate markers in the epithelial cells, while protein expression is more consistently altered in the tumor stroma. The clearly distinctive expression profiles we identified in this study, could potentially be exploited for applications such as image guided resections or potential biomarkers to guide clinical decision making.

## Discussion

The step-wise transition of normal epithelium to CRC has been extensively described from a genetic and cancer cell-centric perspective^2,14^. Several studies have previously defined transcriptomic and epigenetic alterations that accompany tumorigenesis^41–43^, either by comparing normal, adenoma and carcinoma tissues^41,43^ or microdissection of CRC lesions^42^. However, a common limitation of these studies is the inability to spatially contextualize findings. Here, we investigated the transcriptional alterations that accompany CRC onset by spatially-resolved gene expression analysis. Importantly, we defined the biological processes that occur in epithelial and stromal segments separately. This approach revealed pronounced spatially-dependent alterations in oncogenic, metabolic, and immune-related processes during CRC tumorigenesis. An early activation of innate immune responses and expression of innate cytokines was observed. Furthermore, we identified novel biomarker genes that are specifically relevant for the clinical management of early-stage CRC.

The relevance of separate analysis of epithelial and stromal compartments is underlined by specific pathways that showed inversed alterations during the step-wise process of CRC tumorigenesis. For instance, we observed a sharp increase in Notch signaling during tumorigenesis in the stroma, while this pathway decreased in the epithelium. A role for aberrant activation of Notch signaling has previously been implicated in CRC tumorigenesis^44–47^. However, many studies reporting increased expression of Notch and associated genes in human CRC compared to normal tissues are based on bulk transcriptomic data^45,46,48^. The decrease in Notch signaling we observed in the epithelium is most likely a reflection of the important role of Notch signaling in maintenance of normal intestinal epithelium by regulating differentiation of colonic goblet cells and progenitor cells^21,49^. Interestingly, the Wnt-ß catenin pathway has previously been reported to suppress the Notch pathway effector Hath1, resulting in reduced *MUC4* in CRC^50^. The downregulation of the Notch pathway in cancer cells is therefore in line with the reduced expression of the MUC-4 protein that we observed in this study.

The influence of the adaptive immune response on clinically detectable CRCs has been well established^51,52^. Although a large proportion of the immune cells that infiltrate CRCs are of myeloid origin, our understanding of the contribution of the innate immune system on cancer progression is less advanced^53^. Tumor-elicited inflammation is thought to be caused by the breakdown of the epithelial barrier during tumorigenesis that enables microbial products translocate from the intestinal lumen to activate tissue-resident myeloid cells^54,55^. In addition, necrotic cell death as a consequence of hypoxic conditions and lack of sufficient nutrients has been described to induce innate and adaptive immune responses^54^. In this study, we observed an increase in genes associated with innate immunity during the step-wise transition from normal tissue to carcinoma. Already in the transition areas from normal tissue to dysplasia, we detected an upregulation of transcripts for *CFB* and *CCL20* in the epithelium and *CXCL1* in the stroma. Furthermore, we identified an upregulation of major components of the complement pathway, innate cytokine production (IL-1, IL-6, IL-10, IL-18, and TGF-β)^32^, and increase in macrophages in the stroma. These findings suggest an early activation of the innate arm of the immune system during tumorigenesis. The upregulation of *IFITM1* during tumorigenesis, one of our selected biomarkers, also supports this observation, as IFITM proteins have been described to mediate the innate immune role of interferon^56^. Our segmentation approach enabled us to define ligand and corresponding receptor relations of immunomodulator genes. Interestingly, we defined a simultaneous upregulation of *SIRPA* in the stroma and *CD47* in the epithelium during the step-wise progression of CRC. As CD47 expression provides a “don’t eat me”-signal to macrophages expressing SIRPA^38–40^, this could be an effective mechanism of immune escape in early-staged CRCs. Therefore, the CD47/SIRPA axis might also represent an interesting therapeutic candidate for these cancers. Immunotherapeutic strategies that block this innate immune checkpoint are currently in early-phase clinical trials^38,57,58^.

Next to macrophages, we also observed a pronounced increase in estimated fibroblasts and endothelial cells in the stroma during CRC progression. At the same time, our data revealed an increase in enrichment of gene sets reflecting TGF-ß receptor signaling and PDGFRB pathway activation. Both TGF-ß and PDGF are reported to be important growth factors for the conversion of normal fibroblasts into CAFs^16,18,59^. This observation is in concordance with the noticeable increase in expression of collagens in the stromal segment, as CAFs are dominant producers of extracellular matrix proteins^60^. Our results support that the stroma undergoes substantial changes during tumorigenesis. In fact, the alterations within the stroma demonstrated less overall inter-patient variability compared to alterations within the cancer cells themselves. This observation could be relevant for clinical applications that require biomarkers that can discriminate benign from malignant tissue, such as image-guided surgery. Instead of seeking a candidate target that is expressed by the cancer cells, searching for a marker expressed by stromal cells could result in identification of more suitable, universally expressed candidates. Indeed, of the four genes/proteins that were selected because of upregulation in the stroma during CRC development, three (i.e., Thy1, PDGFRB, and Hsp47) strongly correlated with the step-wise progression of CRC in our independent validation cohort.

In addition to gaining fundamental insights in the biological process of CRC tumorigenesis, the spatially-resolved gene expression identified specific biomarkers that could be employed in the clinic. For instance, the identified genes could serve as targets to image-guided endoscopic resections. Importantly, we observed less variability between patients for candidate biomarkers that were expressed in the stroma (i.e., Thy1, PDGFRB, and Hsp47), as opposed to those expressed by cancer cells themselves. This observation is in line the increased heterogeneity between epithelial cells of distinct patients, while stromal cells typically have more shared properties. Future retrospective analyses of larger numbers of pT1 lesions will be necessary to verify whether these same biomarkers can identify those patients that benefit from extensive curative procedures as opposed to a more conservative approach.

Whereas the current study demonstrates the unique biological insights derived by digital spatial profiling of early-stage CRCs, a limitation of the applied approach is the relatively low resolution. Currently, an expanding number of technologies are available to obtain spatially-resolved data^10^. Integration of distinct levels of (single-cell) data obtained from the same biological samples provides unique opportunity to further improve re-define the complex interactions between the cell in the tumor microenvironment during the step-wise progression of CRC.

In conclusion, this work demonstrates the application of spatially resolved gene expression profiling to early-stage CRC and provides insights in the biological processes that accompany the step-wise progression of CRC. Our results indicate an essential role for innate immunity in CRC onset. Furthermore, our approach identified biomarkers that were consistently altered during CRC tumorigenesis. These insights pave the way to improving outcome for early-stage CRC patients.

## Methods

### Human samples

Patient samples were obtained during colonoscopy at the Department of Gastroenterology of the Leiden University Medical Center (LUMC). Patients did not receive any treatment prior to the endoscopic procedure. Endoscopic submucosal dissections (ESDs) were formalin-fixed and paraffine embedded (FFPE). This study was approved by the METC Leiden-Den Haag-Delft (protocol B20.039), and all patients provided informed consent.

### In situ hybridization

To prepare slides for DSP, 5-μm thick FFPE sections of eight different patients (n=8) were deparaffinized, heated in ER2 solution (Leica) at 100 °C for 20 min, and treated with 1 µg/ml proteinase K at 37 °C for 15 min. An overnight in situ hybridization was performed as described with a final probe concentration of 4 nM per probe. Slides were washed twice at 37 °C for 25 min with 50% formamide/2X SSC buffer to remove unbound probes.

### Digital spatial profiling

Prepared slides were incubated with immunofluorescent antibodies and GeoMx Cancer Transcriptome Atlas (CTA, v2.0) profiling reagents simultaneously. Pan-cytokeratin (PanCK; AE1 + AE3, Novus Biologicals, Cat# NBP2-33200AF488) was used for identification of epithelial cells, Vimentin (E-5 clone from Santa Cruz, Cat# sc-373717) for stromal cells, CD45 (D9M8I from CST, Cat# 13917. Internally conjugated) for all hematopoietic cells, and DNA GeoMx Nuclear Stain (Cat# 121303303) for nuclei. Stained slides were loaded onto the GeoMx instrument and scanned.

For each ESD, nine geometric ROIs with different levels of cancer progression were selected by a pathologist, including normal epithelium, transition areas, low-, and high-grade dysplasia, and cancer. Each ROI was UV-illuminated twice, once for the PanCK segment, and once for the Vimentin segment. The resulting areas of illumination (AOIs, n=144) had a surface area in the range of 15,000-350,000 µm^2^, encompassing between 102-5,054 nuclei. GeoMx DSP technology is for research use only and not for use in diagnostic procedures.

### Immunohistochemistry

FFPE tissue sections of 4-μm thick were deparaffinized and rehydrated followed by heat-mediated antigen retrieval using either TE or Citrate buffer depending on primary antibody to be incubated (see Supplementary Table 1). Slides were blocked using SuperBlock (PBS) Blocking Buffer (Thermo Scientific, Cat# 37515) for 30 minutes at RT. Antibodies were diluted in PBS/BSA in predetermined optimal dilutions (see Supplementary Table 1). Tissue sections were incubated overnight at 4°C. Subsequently, slides were incubated with poly-HRP (Thermo Scientific, Cat# 21140) for 60 minutes and developed with DAB chromogen (1:50, DAKO #K3468). Sections were counterstained with hematoxylin (DiaPath #C0305). Slides were dehydrated and mounted with Micromount (Leica #3801731). Consecutive slides were used for the different antibodies. Stained slides were assessed by a pathologist. The intensity of staining was scored across the distinct histological regions for each slide (0 = no expression, 1 = intermediate expression, 2 = strong expression).

### Spatial transcriptomic data processing and normalization

Raw data was available in the GeoMx DSP Analysis Suite. We performed segment and biological probe QC following NanoString’s Cancer Transcriptome Atlas Normalization guidelines. AOI QC was performed in the DSP Analysis Suite to flag low-performing AOIs using the following settings: raw reads threshold <1000 reads, percent aligned reads <80%, sequencing saturation <50%, negative probe count geomean <10, no template PCR control count >60, minimum nuclei <100, and minimum surface area 600 µm^2^. 31 AOIs were flagged with a low negative probe count and one sample with a low sequencing saturation. Probe QC was performed with default settings excluding probes from target count calculation in all segments if (geomean probe in all segments) / (geomean probes withing target) <= 0.1 and if Fails Grubb’s outlier test in >=20% of the segments. If a probe fails Grubb’s outlier test in a specific segment, the probe was excluded from target count calculation in that segment. The limit of quantitation (LOQ) was calculated using 2 standard deviations of the geomean of the negative probes.

To determine whether flagged AOIs need to be removed from the study, we compared background to on-target signal strength. We performed Quartile 3 count (Q3) normalization in the DSP Analysis Suite to account for technical effects between AOIs (e.g., segment area, amount of targetable mRNA, in-situ hybridization binding). While most AOIs have a Q3 signal substantially higher than the NegProbe count, two AOIs needed to be dropped as in these the Q3 signal was comparable to the background. Following quality control (QC), the resulting gene expression matrix consisted of 141 AOIs that passed QC and a total of 1,825 genes. Q3-normalized expression data was exported from the DSP Analysis Suite. All downstream analysis was performed in R (v4.0.3 or later).

### tSNE

Dimension-reduction of the expression matrix was performed using t-Distributed Stochastic Neighbor Embedding by R package “Rtsne” (v.0.15) and visualized using ggplot2 (v.3.3.3).

### Gene set enrichment analysis

Single sample gene set enrichment analysis (ssGSEA) was performed on the log2 transformed, normalized gene expression matrix^61^. Cancer Hallmark pathways were obtained from www.gsea-msigdb.org (H, hallmark gene sets, 50 gene sets, retrieved on 25 February 2021, ^24^). Wiki pathways^28^ were retrieved using the downloadPathwayArchive function of the R package “rWikiPathways” (parameters: organism = “Homo sapiens”, format=“gmt”) on 10 May 2021. All pathways with <2 genes available in the GeoMx CTA probeset were removed.

Downloaded gene sets were tested for applicability for ssGSEA in the CTA expression matrix (1825 genes) by employing the The Cancer Genome Atlas Colon Adenocarcinoma (TCGA-COAD) dataset. ssGSEA was performed on the full TCGA-COAD gene expression matrix and the same matrix filtered to only include the CTA genes. The Pearson correlation between obtained enrichment scores (ES) was determined for each gene set. Gene sets with a Pearson correlation coefficient <0.6, we deemed unsuitable for ssGSEA in our GeoMx CTA dataset and were excluded for further analysis. In the cases that multiple gene sets describe the same biological process or pathway, only the pathway with the highest correlation was maintained. Pathways that describe a specific disease setting or organ were excluded. This resulted in list of 35 Hallmark pathways and 108 WikiPathways.

### Immune cell deconvolution

To estimate relative abundancies from the expression data, we implemented Consensus Tumour Microenvironment cell Estimation (ConsensusTME), a method that relies on integrated gene sets from multiple sources and has been optimized for each individual cancer type, using R package ConsensusTME (v.0.0.1.9)^62^. Parameters were set to “COAD” to specify cancer type and “ssgsea” as statistical method. Corroborating the reliability of this method, we observed low enrichment scores of the immune cell subsets in AOIs from the epithelial compartment (**Extended Data Fig. 3a**).

### Statistical analyses

Differentially expressed genes (DEGs) between PanCK and Vimentin segments were determined using a paired t-test and corresponding False Discovery Rate (FDR) was calculated by the Benjamini-Hochberg method. Stepwise-differential expression analysis between regions with distinct progressive status was calculated using an unpaired t-test. The Spearman correlation test was used to correlate to define genes/pathways that are increasing or decreasing during the stepwise transition of normal colon to carcinoma. For this test, the region parameter was treated as ordinal variable (1= normal; 2 = transition area; 3 = low-grade dysplasia, 4 = high-grade dysplasia, 5 = carcinoma). This analysis was performed for the PanCK and the Vimentin segment separately. Genes with a Spearman correlation >0.5 were mapped to the Global Molecular Network and core network analysis was performed using Ingenuity Pathway Analysis software (IPA). The measurement type to base the analysis was Spearman correlation (Expr Other), and selected Reference Set was Ingenuity Knowledge Base (Genes Only) considering both direct and indirect relationships. Network settings were set to interaction networks with 70 molecules per network.

## Extended Data

**Extended Data Fig. 1.**
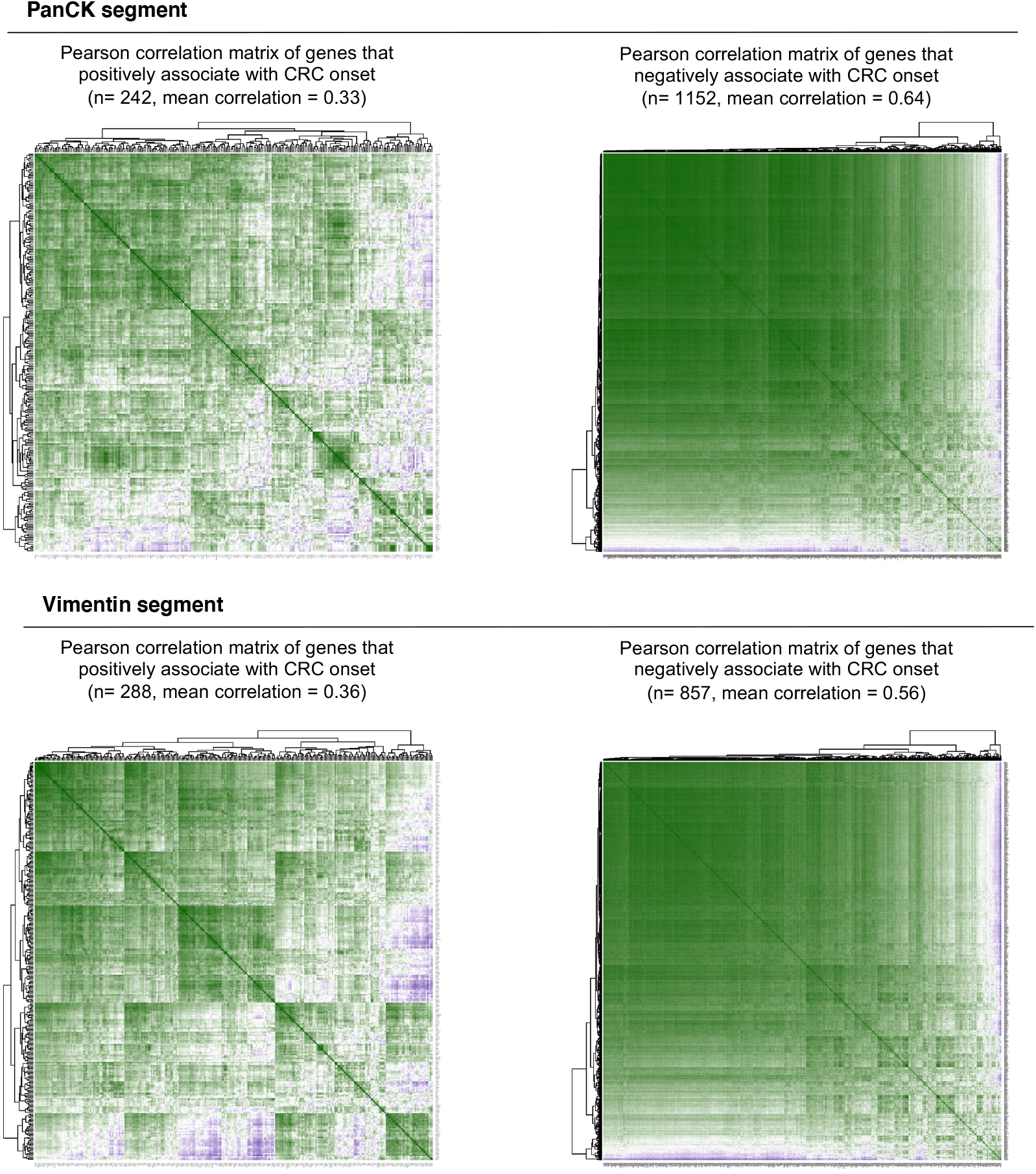
Correlation between genes that associate with CRC tumorigenesis. Spearman correlation matrices between genes that have a significant positive (*left*) or negative (*right*) correlation with CRC tumorigenesis in PanCK (*upper*) or Vimentin (*bottom*) segments. Number of included genes in each category and average correlation is indicated.

**Extended Data Fig. 2.**
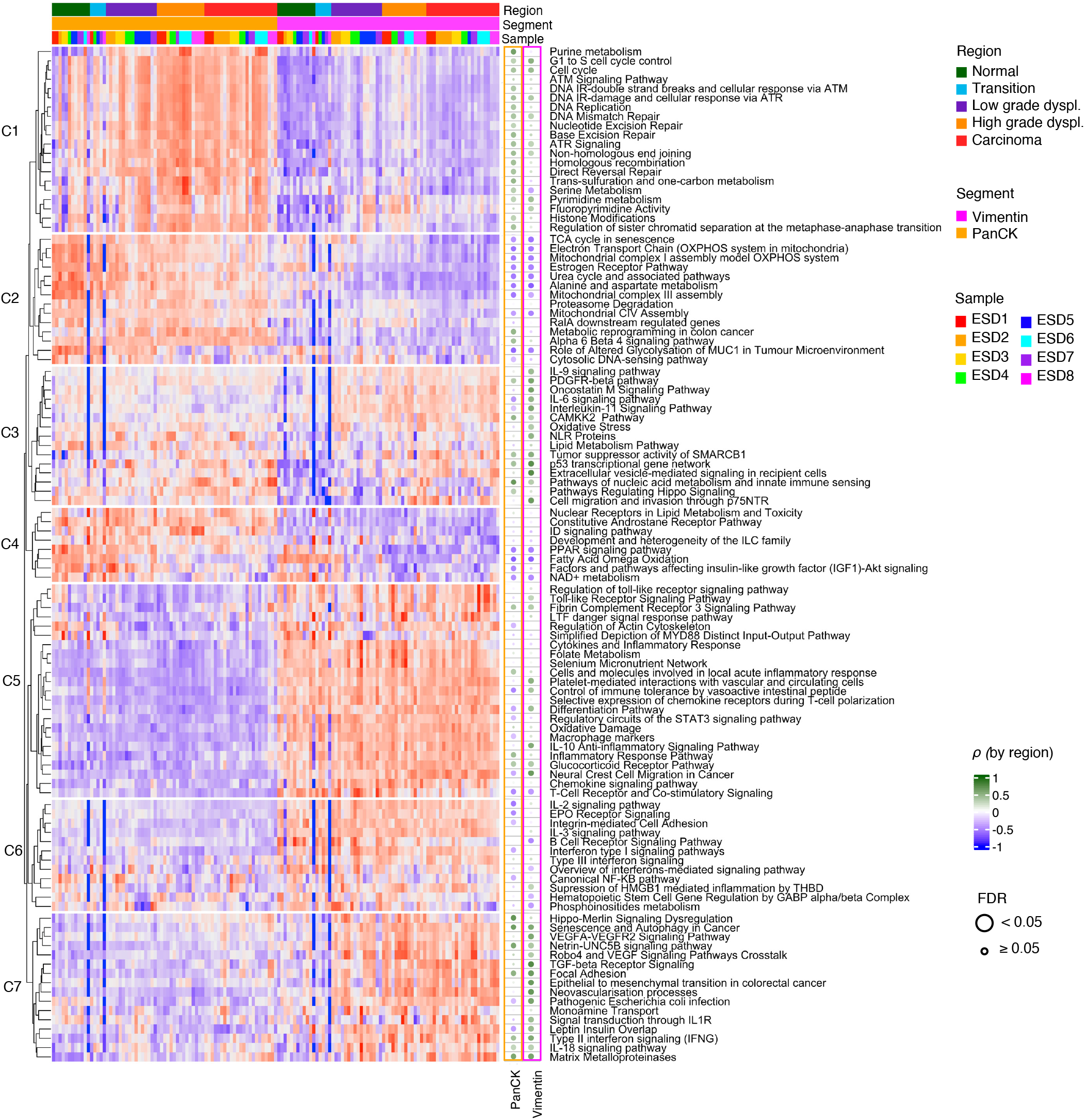
Enrichment of WikiPathway gene sets across distinct AOIs. Unsupervised clustering of gene set enrichment scores calculated using gene sets from the WikiPathways database^28^. All non-redundant biological pathways that were selected for ssGSEA (see methods section) were included in the total collection (n=108 pathways). AOIs are ordered by segment, region, and finally by sample. Main clusters of identified pathways are indicated with white horizontal lines. The corresponding Spearman correlation coefficient, Rho, between histology as ordinal variable and the enrichment scores are indicated for each pathway in PanCK and Vimentin segments separately (dotted heatmap). False Discovery Rate is calculated using Benjamini-Hochberg method. AOI, area of illumination; ssGSEA, single sample gene set enrichment; PanCK, Pan-Cytokeratin; FDR, false discovery rate.

**Extended Data Fig. 3.**
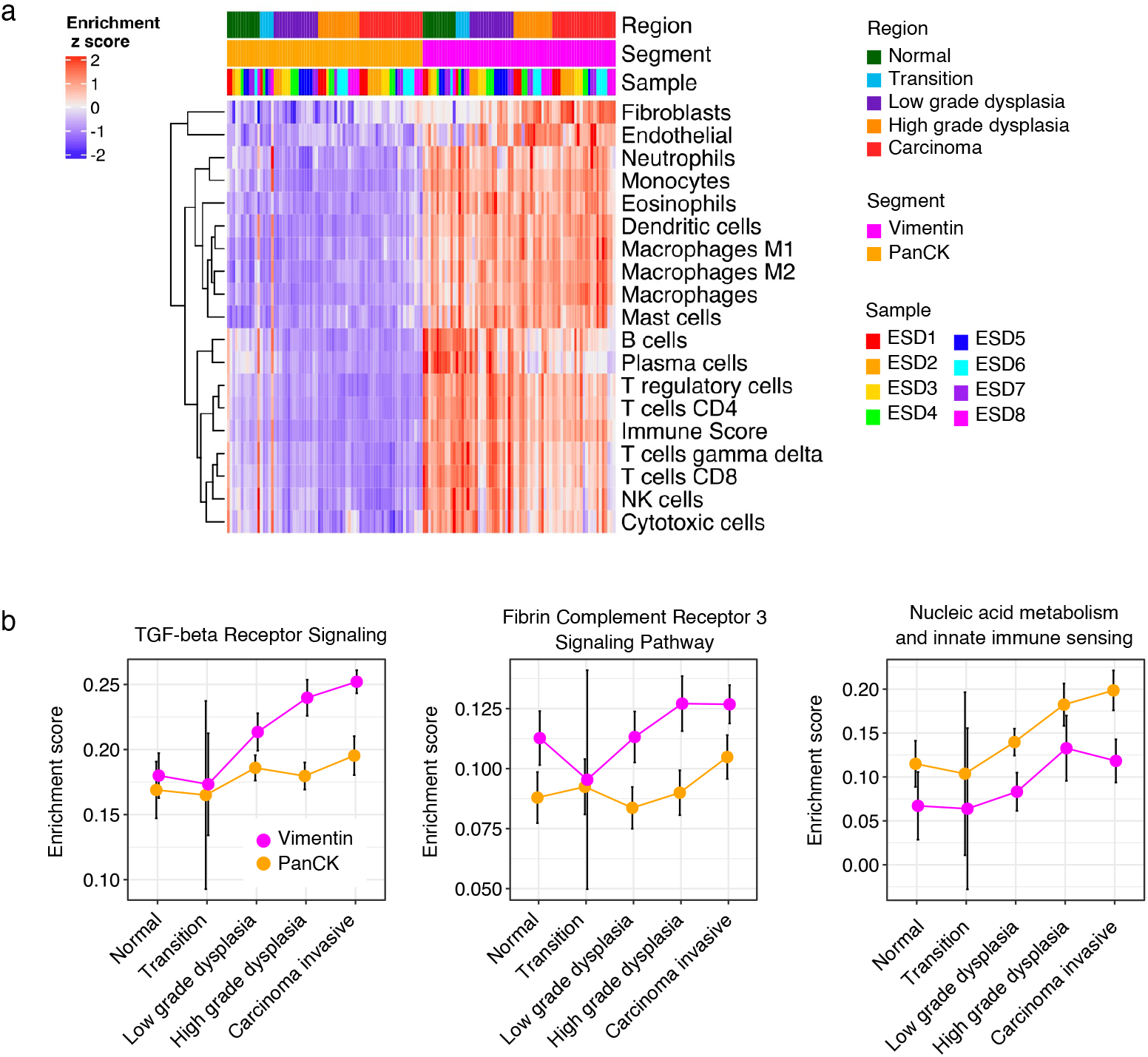
Immune-related alterations by CRC histology. (**a**) Unsupervised clustered heatmap of abundancies of immune cell subsets deconvoluted by the ConsensusTME algorithm^62^. AOIs are ordered by segment, region, and by sample. (**b**) Line graphs of highlighted pathways that increase from normal to carcinoma. Mean enrichment score and corresponding 95%-CI are indicated. CI, confidence interval; PanCK, Pan-Cytokeratin.

## References

1. Fearon, E. R. & Vogelstein, B. A genetic model for colorectal tumorigenesis. Cell 61, 759– 767 (1990).

2. Hanahan, D. & Weinberg, R. A. Hallmarks of Cancer: The Next Generation. Cell 144, 646– 674 (2011).

3. Nowell, P. C. The clonal evolution of tumor cell populations. Science 194, 23–28 (1976).

4. Pickhardt, P. J. et al. Assessment of volumetric growth rates of small colorectal polyps with CT colonography: a longitudinal study of natural history. Lancet Oncol. 14, 711–720 (2013).

5. Kanth, P. & Inadomi, J. M. Screening and prevention of colorectal cancer. BMJ 374, n1855 (2021).

6. Rex, D. K., Shaukat, A. & Wallace, M. B. Optimal Management of Malignant Polyps, From Endoscopic Assessment and Resection to Decisions About Surgery. Clin. Gastroenterol. Hepatol. 17, 1428–1437 (2019).

7. Argilés, G. et al. Localised colon cancer: ESMO Clinical Practice Guidelines for diagnosis, treatment and follow-up. Ann. Oncol. Off. J. Eur. Soc. Med. Oncol. 31, 1291–1305 (2020).

8. Shepherd, N. A. & Griggs, R. K. L. Bowel cancer screening-generated diagnostic conundrum of the century: pseudoinvasion in sigmoid colonic polyps. Mod. Pathol. 28, S88–S94 (2015).

9. Backes, Y. et al. Histologic Factors Associated With Need for Surgery in Patients With Pedunculated T1 Colorectal Carcinomas. Gastroenterology 154, 1647–1659 (2018).

10. de Vries, N. L., Mahfouz, A., Koning, F. & de Miranda, N. F. C. C. Unraveling the Complexity of the Cancer Microenvironment With Multidimensional Genomic and Cytometric Technologies. Front. Oncol. 10, (2020).

11. Merritt, C. R. et al. Multiplex digital spatial profiling of proteins and RNA in fixed tissue. Nat. Biotechnol. 38, 586–599 (2020).

12. Krämer, A., Green, J., Pollard, J. & Tugendreich, S. Causal analysis approaches in Ingenuity Pathway Analysis. Bioinforma. Oxf. Engl. 30, 523–530 (2014).

13. He, T.-C. et al. Identification of c-MYC as a Target of the APC Pathway. Science 281, 1509– 1512 (1998).

14. Vogelstein, B. et al. Cancer genome landscapes. Science 339, 1546–1558 (2013).

15. Hawinkels, L. J. A. C. et al. Active TGF-beta1 correlates with myofibroblasts and malignancy in the colorectal adenoma-carcinoma sequence. Cancer Sci. 100, 663–670 (2009).

16. Calon, A. et al. Dependency of Colorectal Cancer on a TGF-β-Driven Program in Stromal Cells for Metastasis Initiation. Cancer Cell 22, 571–584 (2012).

17. Calon, A. et al. Stromal gene expression defines poor-prognosis subtypes in colorectal cancer. Nat. Genet. 47, 320–329 (2015).

18. Hawinkels, L. J. a. C. et al. Interaction with colon cancer cells hyperactivates TGF-β signaling in cancer-associated fibroblasts. Oncogene 33, 97–107 (2014).

19. Tauriello, D. V. F. et al. TGFβ drives immune evasion in genetically reconstituted colon cancer metastasis. Nature 554, 538–543 (2018).

20. van Es, J. H. et al. Notch/γ-secretase inhibition turns proliferative cells in intestinal crypts and adenomas into goblet cells. Nature 435, 959–963 (2005).

21. Fre, S. et al. Notch signals control the fate of immature progenitor cells in the intestine. Nature 435, 964–968 (2005).

22. Meurette, O. & Mehlen, P. Notch Signaling in the Tumor Microenvironment. Cancer Cell 34, 536–548 (2018).

23. Roumenina, L. T., Daugan, M. V., Petitprez, F., Sautès-Fridman, C. & Fridman, W. H. Context-dependent roles of complement in cancer. Nat. Rev. Cancer 19, 698–715 (2019).

24. Liberzon, A. et al. The Molecular Signatures Database Hallmark Gene Set Collection. Cell Syst. 1, 417–425 (2015).

25. Cantor, J. R. & Sabatini, D. M. Cancer Cell Metabolism: One Hallmark, Many Faces. Cancer Discov. 2, 881–898 (2012).

26. Warburg, O. On respiratory impairment in cancer cells. Science 124, 269–270 (1956).

27. Fernández, L. P., Gómez de Cedrón, M. & Ramírez de Molina, A. Alterations of Lipid Metabolism in Cancer: Implications in Prognosis and Treatment. Front. Oncol. 10, (2020).

28. Slenter, D. N. et al. WikiPathways: a multifaceted pathway database bridging metabolomics to other omics research. Nucleic Acids Res. 46, D661–D667 (2018).

29. Batlle, E. & Massagué, J. Transforming Growth Factor-β Signaling in Immunity and Cancer. Immunity 50, 924–940 (2019).

30. van den Bulk, J., de Miranda, N. F. C. C. & ten Dijke, P. Therapeutic targeting of TGF-β in cancer: hacking a master switch of immune suppression. Clin. Sci. 135, 35–52 (2021).

31. Derynck, R., Turley, S. J. & Akhurst, R. J. TGFβ biology in cancer progression and immunotherapy. Nat. Rev. Clin. Oncol. 18, 9–34 (2021).

32. Lacy, P. & Stow, J. L. Cytokine release from innate immune cells: association with diverse membrane trafficking pathways. Blood 118, 9–18 (2011).

33. Costa, A. et al. Fibroblast Heterogeneity and Immunosuppressive Environment in Human Breast Cancer. Cancer Cell 33, 463–479.e10 (2018).

34. Picarda, E., Ohaegbulam, K. C. & Zang, X. Molecular Pathways: Targeting B7-H3 (CD276) for Human Cancer Immunotherapy. Clin. Cancer Res. Off. J. Am. Assoc. Cancer Res. 22, 3425–3431 (2016).

35. Lee, Y. et al. Inhibition of the B7-H3 immune checkpoint limits tumor growth by enhancing cytotoxic lymphocyte function. Cell Res. 27, 1034–1045 (2017).

36. Oshima, K., Ruhul Amin, A. R. M., Suzuki, A., Hamaguchi, M. & Matsuda, S. SHPS-1, a multifunctional transmembrane glycoprotein. FEBS Lett. 519, 1–7 (2002).

37. Logtenberg, M. E. W., Scheeren, F. A. & Schumacher, T. N. The CD47-SIRPα Immune Checkpoint. Immunity 52, 742–752 (2020).

38. Morrissey, M. A., Kern, N. & Vale, R. D. CD47 Ligation Repositions the Inhibitory Receptor SIRPA to Suppress Integrin Activation and Phagocytosis. Immunity 53, 290–302.e6 (2020).

39. Okazawa, H. et al. Negative regulation of phagocytosis in macrophages by the CD47-SHPS-1 system. J. Immunol. Baltim. Md 1950 174, 2004–2011 (2005).

40. Veillette, A. & Chen, J. SIRPα–CD47 Immune Checkpoint Blockade in Anticancer Therapy. Trends Immunol. 39, 173–184 (2018).

41. Hong, Q. et al. Transcriptomic Analyses of the Adenoma-Carcinoma Sequence Identify Hallmarks Associated With the Onset of Colorectal Cancer. Front. Oncol. 11, 3007 (2021).

42. Qu, X. et al. Integrated genomic analysis of colorectal cancer progression reveals activation of EGFR through demethylation of the EREG promoter. Oncogene 35, 6403–6415 (2016).

43. Maglietta, R. et al. Molecular pathways undergoing dramatic transcriptomic changes during tumor development in the human colon. BMC Cancer 12, 608 (2012).

44. Akiyoshi, T. et al. Gamma-secretase inhibitors enhance taxane-induced mitotic arrest and apoptosis in colon cancer cells. Gastroenterology 134, 131–144 (2008).

45. Jackstadt, R. et al. Epithelial NOTCH Signaling Rewires the Tumor Microenvironment of Colorectal Cancer to Drive Poor-Prognosis Subtypes and Metastasis. Cancer Cell 36, 319–336.e7 (2019).

46. Meng, R. D. et al. γ-Secretase Inhibitors Abrogate Oxaliplatin-Induced Activation of the Notch-1 Signaling Pathway in Colon Cancer Cells Resulting in Enhanced Chemosensitivity. Cancer Res. 69, 573–582 (2009).

47. Sureban, S. M. et al. Knockdown of RNA binding protein musashi-1 leads to tumor regression in vivo. Gastroenterology 134, 1448–1458 (2008).

48. Shaik, J. P. et al. Frequent Activation of Notch Signaling Pathway in Colorectal Cancers and Its Implication in Patient Survival Outcome. J. Oncol. 2020, e6768942 (2020).

49. Riccio, O. et al. Loss of intestinal crypt progenitor cells owing to inactivation of both Notch1 and Notch2 is accompanied by derepression of CDK inhibitors p27Kip1 and p57Kip2. EMBO Rep. 9, 377–383 (2008).

50. Pai, P. et al. MUC4 is negatively regulated through the Wnt/β-catenin pathway via the Notch effector Hath1 in colorectal cancer. Genes Cancer 7, 154–168 (2016).

51. Galon, J. et al. Type, density, and location of immune cells within human colorectal tumors predict clinical outcome. Science 313, 1960–1964 (2006).

52. Pagès, F., Galon, J. & Fridman, W. H. The essential role of the in situ immune reaction in human colorectal cancer. J. Leukoc. Biol. 84, 981–987 (2008).

53. Kather, J. N. & Halama, N. Harnessing the innate immune system and local immunological microenvironment to treat colorectal cancer. Br. J. Cancer 120, 871–882 (2019).

54. Schmitt, M. & Greten, F. R. The inflammatory pathogenesis of colorectal cancer. Nat. Rev. Immunol. 1–15 (2021) doi:10.1038/s41577-021-00534-x.

55. Grivennikov, S. I. et al. Adenoma-linked barrier defects and microbial products drive IL-23/IL-17-mediated tumour growth. Nature 491, 254–258 (2012).

56. Brass, A. L. et al. The IFITM Proteins Mediate Cellular Resistance to Influenza A H1N1 Virus, West Nile Virus, and Dengue Virus. Cell 139, 1243–1254 (2009).

57. Jalil, A. R., Andrechak, J. C. & Discher, D. E. Macrophage checkpoint blockade: results from initial clinical trials, binding analyses, and CD47-SIRPα structure–function. Antib. Ther. 3, 80–94 (2020).

58. Matlung, H. L., Szilagyi, K., Barclay, N. A. & van den Berg, T. K. The CD47-SIRPα signaling axis as an innate immune checkpoint in cancer. Immunol. Rev. 276, 145–164 (2017).

59. Bronzert, D. A. et al. Synthesis and secretion of platelet-derived growth factor by human breast cancer cell lines. Proc. Natl. Acad. Sci. 84, 5763–5767 (1987).

60. Kalluri, R. The biology and function of fibroblasts in cancer. Nat. Rev. Cancer 16, 582–598 (2016).

61. Barbie, D. A. et al. Systematic RNA interference reveals that oncogenic KRAS-driven cancers require TBK1. Nature 462, 108–112 (2009).

62. Jiménez-Sánchez, A., Cast, O. & Miller, M. L. Comprehensive Benchmarking and Integration of Tumor Microenvironment Cell Estimation Methods. Cancer Res. 79, 6238– 6246 (2019).

